# Sex-specific age-related changes in glymphatic function assessed by resting-state functional magnetic resonance imaging

**DOI:** 10.1101/2023.04.02.535258

**Authors:** Feng Han, Xufu Liu, Yifan Yang, Xiao Liu

## Abstract

The glymphatic system that clears out brain wastes, such as amyloid-β (Aβ) and tau, through cerebrospinal fluid (CSF) flow may play an important role in aging and dementias. However, a lack of non-invasive tools to assess the glymphatic function in humans hindered the understanding of the glymphatic changes in healthy aging. The global infra-slow (<0.1 Hz) brain activity measured by the global mean resting-state fMRI signal (gBOLD) was recently found to be coupled by large CSF movements. This coupling has been used to measure the glymphatic process and found to correlate with various pathologies of Alzheimer’s disease (AD), including Aβ pathology. Using resting-state fMRI data from a large group of 719 healthy aging participants, we examined the sex-specific changes of the gBOLD-CSF coupling, as a measure of glymphatic function, over a wide age range between 36-100 years old. We found that this coupling index remains stable before around age 55 and then starts to decline afterward, particularly in females. Menopause may contribute to the accelerated decline in females.

## Introduction

Aging is the leading risk factor for cognitive decline and neurodegenerative disorders that are often associated with excessive accumulation of misfolded proteins, including the amyloid-β and tau, in the brain (Hou et al., 2019; Mawuenyega et al., 2010; Peng et al., 2016). Recent studies suggested the aggregation of toxic proteins could be partly attributed to impaired glymphatic clearance at advancing ages (Benveniste et al., 2019; Boland et al., 2018; Jessen et al., 2015). The glymphatic system, as “glia-lymphatic system”, constitutes a pathway for brain waste clearance in the central nervous system (Iliff et al., 2012; Jessen et al., 2015). In this clearance pathway, cerebrospinal fluid (CSF) moves from the periarterial space, facilitated by aquaporin-4 (AQP4) channels in astroglial endfeet, into the interstitial space to wash out interstitial solutes, including Aβ and tau, into the perivenous space surrounding deep-draining veins (Jessen et al., 2015; Tarasoff-Conway et al., 2015). The paravascular CSF recirculation and interstitial solute efflux have been found to decrease in aged mice due to widespread loss of perivascular AQP4 polarization and reduced pulsatility of intracortical arterioles (Kress et al., 2014). In humans, the clearance along the glymphatic pathway and downstream meningeal lymphatic vessels, measured by contrast-agent MRI, decreased and delayed in older patients as compared with younger ones (Zhou et al., 2020). However, the invasive nature of these imaging tools has hindered a large-scale study of glymphatic function in healthy aging subjects. As a result, it remains unclear how the glymphatic function changes in aging, which is vital to understanding the mechanisms of age-related neurodegenerative disorders and cognitive decline.

Global infra-slow (< 0.1 Hz) brain activity measured by resting-state fMRI (rsfMRI) was recently linked to the glymphatic function (Kiviniemi et al., 2016) and used to quantify its changes in Alzheimer’s disease (AD) (Han et al., 2021b) and Parkinson’s disease (PD) patients (Han et al., 2021a). Increasing evidence suggested the brain exhibits highly structured, brain-wide infra-slow (< 0.1 Hz) activity during the resting state (Gu et al., 2021; Liu et al., 2021; Raut et al., 2021; Thompson et al., 2014). This global brain activity is evident in neural signals of distinct scales, ranging from single neuron recordings to whole-brain fMRI, and closely related to transient arousal modulations (Gu et al., 2021; Liu et al., 2021). The fMRI measure of this activity, i.e., the global mean rsfMRI blood-oxygenation-level-dependent (gBOLD) signal, is coupled by large CSF movements (Fultz et al., 2019) and astroglial calcium waves (Wang et al., 2018), suggesting its potential link to the glymphatic function. The gBOLD is greatly enhanced during sleep (Fukunaga et al., 2006; Larson-Prior et al., 2009; Olbrich et al., 2009), in accordance with the sleep-enhanced nature of the glymphatic function (Xie et al., 2013). In contrast, arterial and respiratory pulsations, which had been traditionally regarded as the main glymphatic drivers (Iliff et al., 2013; Yamada et al., 2013), actually lack of this attribute with the decreased amplitude during sleep (Boudreau et al., 2013; Douglas et al., 1982; Snyder et al., 1964). For all these reasons, the strength of gBOLD-CSF coupling has been proposed as a surrogate measure of the glymphatic function and found to correlate with various AD pathologies and cognitive decline in PD (Han et al., 2021b, 2021a). Recently, the disengagement of gBOLD from the default mode network (DMN) regions were found to account for early, preferential Aβ accumulation in these higher-order brain areas in the early stage of Aβ pathology (Han et al., 2022). In these early studies, the gBOLD-CSF coupling also was found to correlate significantly with age and sex (Han et al., 2021b, 2021a), but the related findings were limited by narrow age ranges and complicated by the inclusion of patient data.

The fMRI-based glymphatic measurement, together with the widely available rsfMRI data, provides us a unique opportunity to study the change of the glymphatic function over a wider age range and in a larger population of healthy aging subjects. In this study, we used rsfMRI data of 719 healthy aging subjects in the Human Connectome Project Aging (HCP-A) (Harms et al., 2018) to study the age-related changes in glymphatic function in a sex-specific way. We found that the fMRI-based glymphatic measure remained relatively stable within the range of age 36-54 and then started to decrease around age mid-50s. Compared with males, females showed a larger and more abrupt decline of the glymphatic function at this transitioning point. In addition, menopause may lead to an accelerated glymphatic decline in females.

## Results

### Nonlinear age trajectory of the gBOLD-CSF coupling

We used rsfMRI data of 719 healthy subjects (403 females) from the HCP-A project with age ranging from 36 to 100. Following previous procedures (Fultz et al., 2019; Han et al., 2021b, 2021a), we obtained the whole-brain gBOLD signal from the gray matter regions and the bottom-slice CSF signal around the bottom of the cerebellum to measure CSF movements via MR inflow effects (Fultz et al., 2019; Gao et al., 1996; Gao and Liu, 2012) (**Fig. S1A** and **S1B**). Consistent with the previous studies, the averaged cross-correlation function of the two signals displayed a biphasic pattern with a negative peak (*r* = -0.33) at the +3.2 sec lag (**Fig. S1C**). The gBOLD-CSF correlation at this +3.2 sec time lag was then computed for individual subjects to quantify their coupling and thus the glymphatic function. The gBOLD-CSF coupling was then averaged within 10 equal-size groups of subjects at different ages, and the resulting age-related trend displayed a clear non-linear trajectory: it remains relatively stable between ages 36 to 54 and then begins to decline at around 55 years old (i.e., yrs) (**Fig. 1A**). We then divided the entire cohort into younger and older groups according to the age boundary (53.9 yrs) between the fourth and fifth groups, which had the largest drop among pairs of consecutive groups. The gBOLD-CSF coupling is not correlated (*r* = -3.7×10^-3^, *p* = 0.95) with age in the younger group whereas this correlation is significant (*r* = 0.14, *p* = 3.9×10^-3^) in the older group (**Fig. 1B**).

**Fig. 1.**
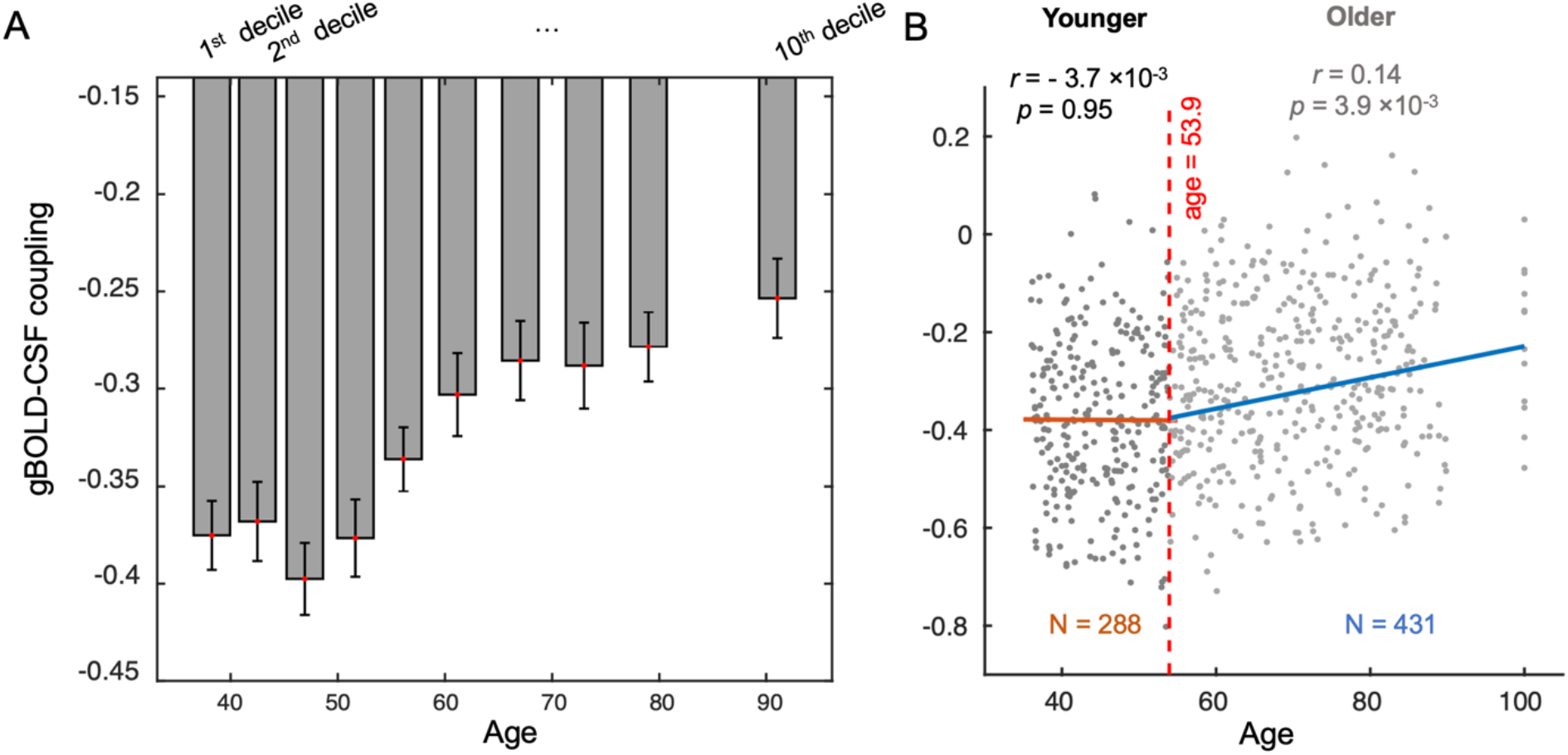
The change of gBOLD–CSF coupling with age. (**A**) The gBOLD–CSF coupling remains relatively stable before age mid-50s and then begins to decrease since then. The entire cohort was grouped into 10 subgroups of equal size. Each error bar represents one standard error of the mean. (**B**) The gBOLD-CSF coupling was significantly correlated with age for the group of subjects over 53.9 yrs (*r* = 0.14, *p* = 3.9×10^-3^; gray dots at the right), but not so (*r* = -3.7×10^-3^, *p* = 0.95; black dots at the left) for the younger group. Each dot represents one subject.

We examined potential factors mediating this age-coupling association. Neither the coupling index nor age was significantly correlated with the total Pittsburgh Sleep Quality Index (PSQI) score, but they were similarly correlated with a few PSQI items, including sleep medication, trouble sleeping, and sleep hours. Nevertheless, the age-related changes in the gBOLD-CSF coupling remain largely unchanged with adjusting for these sleep-related measurements (**Fig. 2**). Likewise, the age trajectory of the gBOLD-CSF remained similar with controlling for other non-sleep factors, including the head motion assessed by mean framewise displacement (FD) (**Fig. S2**), the brain volume (**Fig. S3**), and the CSF volume (mainly from the ventricles) (**Fig. S4**), even though these factors showed a significant dependence on age (all *p* < 3.9×10^-7^ for linear regression).

**Fig. 2.**
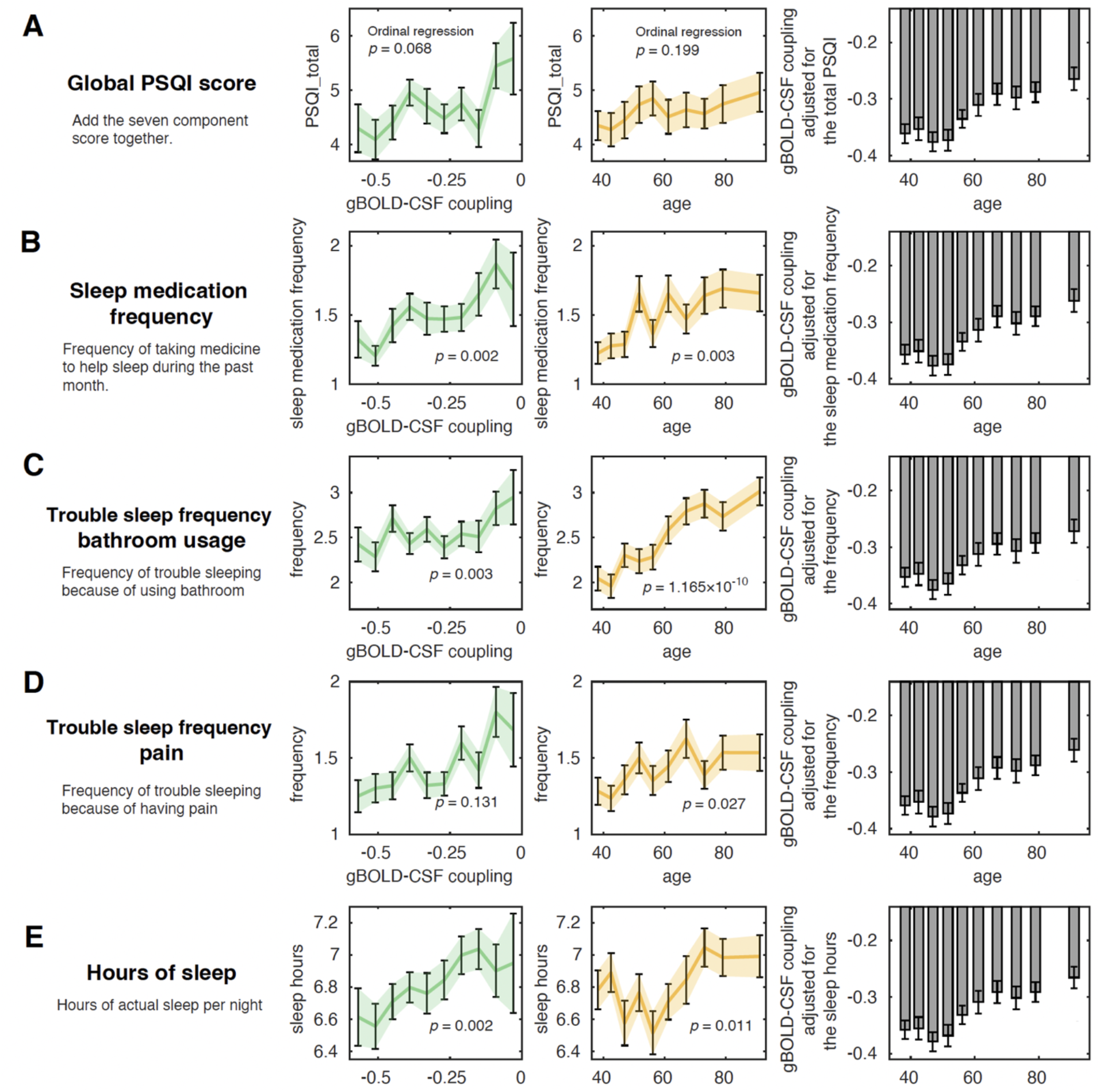
The age-related changes in gBOLD-CSF coupling remain similar with controlling for sleep measures. The gBOLD-CSF coupling (left) and age (middle) are similarly correlated with a few sleep-related measurements in PSQI, including the sleep medication frequency (2^nd^ row), the frequency of bathroom-use during sleep-time (3^rd^ row), the frequency of having pain during sleep-time (4^th^ row), and sleep hours (5^th^ row), but not the total PSQI score (1^st^ row). However, the age-related changes in gBOLD-CSF coupling remain similar with adjusting for these PSQI measures (right). Each dot represents one subject, and error bars represent the standard error of the mean.

### Sex-specific differences in age-related changes

The gBOLD–CSF coupling is different between females and males. The mean gBOLD-CSF strength is significantly (*p* = 7.9×10^-6^, two-sample t-test) weaker in females than males (**Fig. 3A**), consistent with the previous finding in an AD cohort (Han et al., 2021b). The age trajectories of the coupling index are different for the two groups. The gBOLD–CSF coupling decreases more evidently and abruptly with aging in the females, particularly around 55 yrs, whereas the males showed a more gradual and steady age-related changes (**Fig. 3B**).

**Fig. 3.**
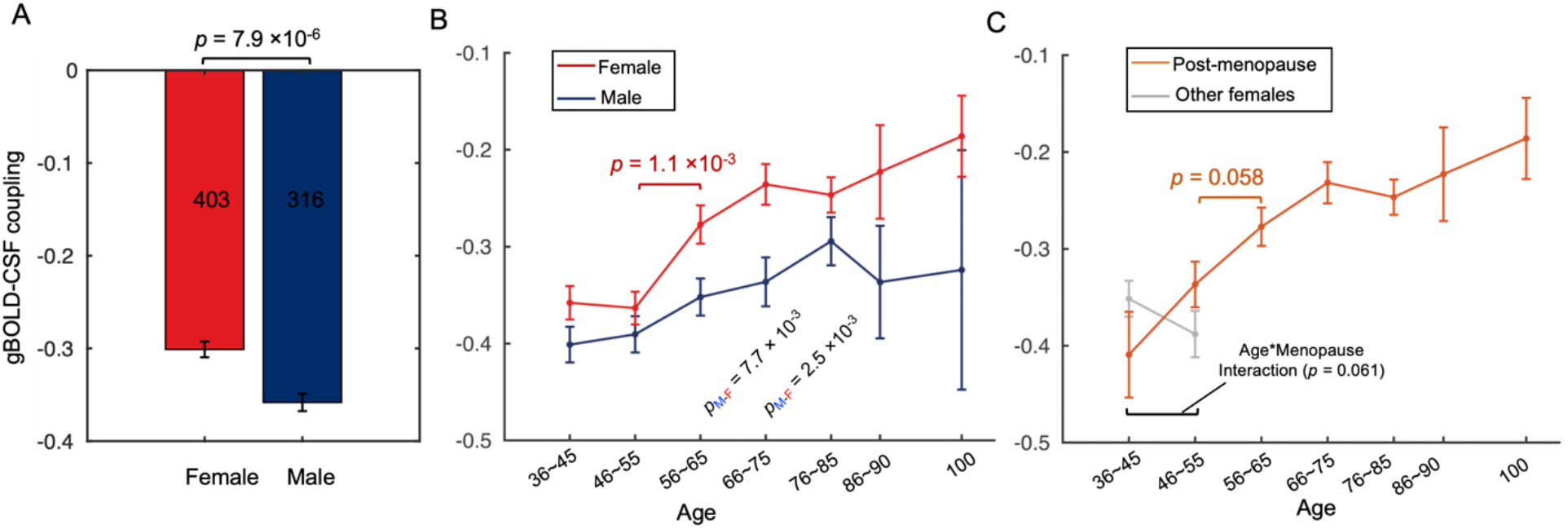
The age-coupling association showed distinct patterns between males and females. (**A**) The gBOLD-CSF coupling is significantly weaker in females (red) than in males (blue) (*p* = 7.9×10^-6^, two-sample t-test). (**B**) The age-related changes of the gBOLD-CSF coupling were summarized separately for the females and males and showed different patterns of trajectory: gBOLD–CSF coupling in the females showed a steep decline around 55 yrs (*p* = 1.1×10^-3^ for two consecutive subgroups around that age), in contrast to the slow and gradual decreases in the males. (**C**) The gBOLD-CSF coupling trends in the “Post-menopause” and “Other females” groups. All females above age 55 are post-menopause. Within the age range of 36-55, the menopause status and age showed a marginally significant (*p* = 0.061) interaction effect on the gBOLD-CSF coupling. The coupling index started to decrease early at age mid-40s and thus its reduction around age mid-50s is less significant (*p* = 0.058) as compared with the entire female group.

We further tested whether and how the gBOLD-CSF coupling is affected by menopause, which often occurs before age 55. This analysis was focused on the age range of 36-55, which includes both post-menopause and other females. The menopause status had no significant effects on the gBOLD-CSF coupling strength for two subgroups of subjects of similar ages (both *p* > 0.13; two sample t-test). However, it showed a marginally significant (*p* = 0.061) interaction with age on the coupling strength. Consistent with this result, the gBOLD-CSF coupling in the postmenopausal females appeared to decline earlier in the age mid-40s (**Fig. 3C**), making the drop around age 55 less abrupt and significant (*p* = 0.058). The result suggests the potential effect of the menopause to accelerate the decline of the fMRI-based glymphatic measure.

## Discussion

Here we study the sex-specific age-related change in the glymphatic function in a large healthy aging population based on the gBOLD-CSF coupling measured by rsfMRI. We showed that the gBOLD-CSF coupling changes with age in a non-linear way: it remains relatively stable from 36 yrs to 54 yrs and then begins to decrease afterwards. Importantly, the decrease at age mid-50s is larger and more abrupt in females than in males. Converging evidence has suggested a link between the glymphatic function and the resting-state global brain activity, often measured by the gBOLD signal of rsfMRI. The gBOLD, once regarded as noise, measured a brain-wide, low-frequency (< 0.1 Hz) activity linked to transient arousal modulations, which also has been observed in monkey electrocorticography (ECoG) (Liu et al., 2015, 2018) and mice spiking data as highly structured patterns (Liu et al., 2021). The low-frequency rsfMRI signals were first linked to the glymphatic function due to their potential link to CSF dynamics and vasomotor waves (Kiviniemi et al., 2016). It was found later that it is the global component of rsfMRI, i.e., gBOLD, that is coupled to CSF movements in a sleep-dependent way (Fukunaga et al., 2006; Fultz et al., 2019; Helakari et al., 2022; Larson-Prior et al., 2009; Olbrich et al., 2009). The sleep dependency makes it more suitable for driving the sleep-dependent glymphatic clearance (Holth et al., 2019; Xie et al., 2013), as compared with the cardiac and respiratory pulsations that are actually suppressed during sleep (Baust and Bohnert, 1969; Boudreau et al., 2013; Douglas et al., 1982; Guazzi and Zanchetti, 1965; Snyder et al., 1964). Nevertheless, the gBOLD is not independent from these physiological drivers but shows strong correlations with the low-frequency modulation of cardiac and respiratory functions (Birn et al., 2006; Chang et al., 2009; Gu et al., 2020; Özbay et al., 2019, 2018; Power et al., 2018). Consistent with these human findings of strong low-frequency physiological modulation, a recent mice study demonstrated very strong arterial constrictions/dilations during sleep in the same frequency range (<0.1 Hz) (Turner et al., 2020). Such low-frequency vessel modulations were coupled by pupil size changes suggestive of transient arousal modulations (Turner et al., 2022), similar to the global brain activity measured by gBOLD (Liu et al., 2021, 2018; Pais-Roldán et al., 2020; Turchi et al., 2018). Animal research also suggested that the gBOLD is coupled by large calcium signals of astrocytes (Pais-Roldán et al., 2020), and the AQP4 channels on the endfeet of these cells are a key player of the glymphatic system (Iliff et al., 2012; Jessen et al., 2015). To date, the key evidence linking the gBOLD to the glymphatic function came from a human study showing that the gBOLD is coupled to large CSF movements (Fultz et al., 2019). Based on this, the gBOLD-CSF coupling was used to quantify glymphatic function and found correlated with various pathologies of AD and cognitive decline in PD (Han et al., 2021b, 2021a). The preferential reduction of gBOLD signal in the higher-order default mode network was found to account for early, preferential β-amyloid accumulations in the same regions at the early stage of AD (Han et al., 2022). All these findings established the foundation for using the gBOLD-CSF coupling to measure the glymphatic function. But a direct proof of their relationship would need future experiments capable of recording brain signals across distinct spatial and temporal scales. It is also worth noting that the debate is ongoing regarding specific components of glymphatic theory, e.g., the convective flow in the interstitial space and the involvement of AQP4 channels in the process (Abbott et al., 2018; Hladky and Barrand, 2022). Nevertheless, a consensus view is reached regarding the existence of periarterial CSF flow (Mestre et al., 2018) and its role in waste clearance (van Veluw et al., 2020), which is more related to the gBOLD-CSF coupling in the present study.

Brain aging affects the glymphatic function. The CSF inflow of larger tracers or macromoleculars decreased up to 85% in old wild-type mice as compared to young counterpart (Da Mesquita et al., 2018; Nedergaard and Goldman, 2020). The decreased glymphatic flow in aged mice has been partly attributed to dysregulation of astroglial water transport due to the widespread loss of AQP4 polarization (Kress et al., 2014), the decline of CSF pressure (Fleischman et al., 2012), and the changes of CSF secretion and protein content (Chen et al., 2009). Arterial wall stiffening and associated reduction of arterial pulsatility (Zieman et al., 2005) may also account for age-related glymphatic reduction (Iliff et al., 2013). Human studies of glymphatic function remain sparse (Eide et al., 2018; Zhou et al., 2020). Previous studies using the gBOLD-CSF coupling found a consistent association between the age and glymphatic function, but only in patients of relatively old ages (Han et al., 2021a, 2021b). A retrospective study combined contrast-agent MRI data from various patient groups to study the glymphatic function and its change with age (Zhou et al., 2020). Despite different methodologies and patient cohorts from the present study, a similar age trajectory was observed for the glymphatic function: it remains stable before 50 yrs and then begins to decline since then (Zhou et al., 2020). The age-related glymphatic changes could be critical for the aging-related risk for neurodegenerative diseases (Hou et al., 2019). The glymphatic dysfunction may result in inadequate clearance and thus accumulation of toxic proteins, such as Aβ and tau, and thereby increase the vulnerability to developing cognitive impairments and neurodegenerative diseases (Jessen et al., 2015; Tarasoff-Conway et al., 2015). Epidemiologic research suggested the late-onset AD, the most common AD variant, starts around the mid-60s with the prevalence doubled every 5 years afterwards (Qiu et al., 2009). But the pathophysiological process, including the accumulation of aggregated Aβ, could begin more than a decade before the dementia (Jack et al., 2013; Sperling et al., 2014). Together, these findings suggested a timeline consistent with our finding that the glymphatic function begins to decrease at age mid-50s.

The female sex is another leading risk factor for developing AD (Mielke et al., 2014). It has been found a significantly weaker glymphatic function, as measured by gBOLD-CSF coupling strength, in females than in males (Han et al., 2021b). Here we confirmed the finding with a much larger dataset from healthy populations. Importantly, we further showed that the glymphatic function had different age trajectories in the two groups with the females displaying a larger and more abrupt decline at around 55 yrs. In fact, women indeed show larger and faster cognitive declines than men with aging (Levine et al., 2021; Nooyens et al., 2022). Together with increasing evidence that links the glymphatic dysfunction and cognitive impairments (Da Mesquita et al., 2018; Iliff et al., 2014; Zou et al., 2019), the sex-specific glymphatic change with aging may provide a possible explanation for the sex differences in age-related cognitive decline. The menopause and associated hormone loss have been suggested to contribute to cognitive decline in females (Brown and Gervais, 2020; Hachul et al., 2015). Our result is not inconsistent with this notion by showing a marginally significant (*p* = 0.061) interaction between age and menopause on the coupling metrics. Among the females of age 36-55, the post-menopausal group appeared to show a decline of gBOLD-CSF coupling with age, which is absent in the non-menopause group. A limited sample size and information related to menopause could partially account for statistical non-significance. But the finding should warrant future studies looking into the menopause effects on the glymphatic function with a refined design and/or augmented dataset.

Sleep quality appeared not to be a major contributor to the age-related glymphatic changes. The gBOLD-CSF coupling and age are similarly correlated with a few sleep measures, including the frequency of using sleep medication and the sleep troubles in the month prior to the experiments. However, the age-related changes in the gBOLD-CSF coupling remained largely unchanged after adjusting for these sleep measures. Nevertheless, the change in sleep architecture might be related to the sex difference seen in the age-related glymphatic changes. The age and sex are known to have a strong interaction effect on the composition of sleep stages. Aging in males is associated with a significant increase of light sleep (stages 1&2) but decrease in slow wave sleep (SWS: sleep stages 3&4), whereas this age-related change is absent in women (Mander et al., 2017; Redline et al., 2004). It is known that the glymphatic function increases during sleep and anesthesia featuring strong slow wave activity (SWA) (Hablitz et al., 2019; Ju et al., 2017; Xie et al., 2013), and one would thus expect an improved glymphatic clearance with a higher percentage of SWS. The empirical evidence, however, suggested an opposite by showing higher SWS is associated with lower CSF Aβ42 level (Varga et al., 2016), which is an early indicator of preclinical AD and often accompanied by cortical Aβ accumulation (Jack et al., 2013; Palmqvist et al., 2017). The paradox might be explained by an observation about the gBOLD and its coupling with CSF flow. They were more specifically related to ultra-slow (0.6-1 Hz) component of SWA (often related to K-complexes) (Özbay et al., 2019) and phasic changes of SWA power (Fultz et al., 2019; Gu et al., 2022), which could be stronger during the light sleep than SWS. Indeed, the SWA was found to decrease in subjects with more cortical Aβ and poorer memory consolidation (Mander et al., 2015; Winer et al., 2020), as well as AD patients (De Gennaro et al., 2017), but the reduction was specific to its ultra-slow component (0.6-1 Hz) with the delta-band (1-4 Hz) power showing opposite changes. Based on all these findings, it is possible that the age-related increase in the percentage of light sleep in males may help to offset some age-related decline in glymphatic function and thus lead to its slow deterioration as compared with females. However, the test of this hypothesis would have to be left for future studies, particularly those with assessment of subjects’ sleep architecture.

## STAR Methods

### Participants and study data

This study included 719 healthy human subjects (36∼100 yrs; 403 females) who have participated in all 4 sessions of rsfMRI scanning in the HCP-A project (https://www.humanconnectome.org/study/hcp-lifespan-aging). For these subjects, we also used their T1-weighted structural MRI imaging and demographic data, such as the age, sex, and the menstrual cycle of females. These “typical aging” subjects were healthy for their age without identified pathological causes of cognitive declines, such as stroke or clinical dementia (Bookheimer et al., 2019). All participants provided written informed consent, and investigators at each HCP-A participating site obtained ethical approval from the corresponding institutional review board (IRB).

The use of de-identified data from the HCP-A and the sharing of analysis results have been reviewed and approved by the Pennsylvania State University IRB (IRB#: STUDY00008766) and also strictly followed the National Institute of Mental Health (NIMH) Data Archive-data use certification (DUC).

### Image acquisition and preprocessing

The rsfMRI data were acquired at 3T MR scanners (Siemens Medical Solutions; Siemens, Erlangen, Germany) with a matched protocol (Harms et al., 2018) across four acquisition sites including Washington University St. Louis, University of Minnesota, Massachusetts General Hospital, and University of California, Los Angeles (researchers in Oxford University dedicating to the data analysis). For each subject, 4 sessions of rsfMRI (including the anterior to posterior phase encoding (PE) from Day1, i.e., AP1, as well as PA1, AP2, and PA2) were followed by one T1-weighted structural MRI session (MPRAGE sequence, echo time (TE)= 1.8/3.6/5.4/7.2 ms [multi-echo], repetition time (TR) = 2,500 ms, field of view (FOV) = 256 × 256 mm^2^, 320 × 300 matrix, number of slices = 208, voxel size = 0.8 × 0.8 × 0.8 mm^3^, flip angle = 8°) (Harms et al., 2018). The T1-weighted MRI served to provide the whole brain and CSF volume information and was used for the anatomical segmentation and registration. For rsfMRI acquisition, 488 fMRI volumes were collected with a multiband gradient-recalled (GRE) echo-planar image (EPI) sequence (TR/TE=800/37 ms, flip angle=52°, FOV = 208 mm, 104 × 90 matrices, 72 oblique axial slices, 2 mm isotropic voxels, multiband acceleration factor of 8).

We referred to the previous study (Han et al., 2021b) in preprocessing the rsfMRI data using the FSL (version 5.0.9; https://fsl.fmrib.ox.ac.uk/fsl/fslwiki/FSL) (Smith et al., 2004) and AFNI (version 16.3.05; https://afni.nimh.nih.gov/) (Cox, 1996). The general fMRI preprocessing procedures included motion correction, skull stripping, spatial smoothing (full width at half maximum (FWHM) = 4mm), temporal filtering (bandpass, approximately 0.01 to 0.1 Hz), and the co-registration of each fMRI volume to corresponding T1-weighted structural MRI and then to the 152-brain Montreal Neurological Institute (MNI-152) space. The motion parameters were not regressed out to avoid attenuating the gBOLD signal (Gu et al., 2020; Han et al., 2021b). The preprocessing of structural images was performed using FSL. Processing steps included spatial normalization and skull stripping.

### Extract gBOLD and the CSF inflow signals and compute their coupling

We followed the previous studies (Han et al., 2021b, 2021a) to extract the gBOLD signal and CSF inflow signal. We defined the mask of the gray matter regions based on the Harvard-Oxford cortical and subcortical structural atlases (https://neurovault.org/collections/262/). We then transformed the gray-matter mask from the MNI-152 space back to the original space of each session to avoid spatial blurring from the registration process (Fultz et al., 2019), and spatially averaged the Z-normalized gray-matter rsfMRI signals to obtain the gBOLD signal. Following the previous study (Han et al., 2021b), the CSF inflow signal was extracted from the CSF region at the bottom edge of fMRI, with a similar voxel number of CSF ROI for all subjects/sessions (see an example in **Fig. S1A**). We extracted the CSF ROI from the preprocessed fMRI signal at the original individual space referring to the corresponding CSF region below the bottom of the cerebellum from the high-resolution T1-weighted MRI (see an exemplary time-series of gBOLD and bottom CSF fMRI in **Fig. S1B**).

The cross-correlation function was calculated on the extracted gBOLD signal and the CSF inflow signal for each fMRI session from each individual subject. The cross-correlation function quantified the Pearson’s correlation at different time lags. We first averaged all the cross-correlation functions from the 4 individual fMRI sessions (AP1, PA1, AP2, and PA2) for each subject, and further averaged the functions across all subjects (**Fig. S1C**). Referring to the previous studies (Han et al., 2021b, 2021a), we quantified the gBOLD–CSF coupling with the session-mean cross-correlation at the lag of +3.2 seconds, where the negative peak of the subject-mean cross-correlation located, for each subject.

### The relationship between the gBOLD–CSF coupling and age

To access the association between the gBOLD–CSF coupling and age, we first divided all the subjects into 10 sub-groups with different ages (based on the deciles), and further calculated and compared the mean gBOLD–CSF coupling for each sub-group. Moreover, all the subjects were separated into the younger (age < 53.9 yrs) and older (age ≥ 53.9 yrs) groups. The linear regression was used to evaluate the association between the ages and the coupling measures for the subjects in each sub-group.

Several sensitivity analyses were then performed to test whether the age-coupling association would be driven by the various sleep quality measures (i.e., PSQI measures), the brain volume, as well as the CSF volume. We included the total PSQI score, as well as 4 different PSQI items, which were selected due to their strong dependence (*p* < 0.05 for linear regression or ordinal regression) with the coupling measures or age, covering the components/aspects of sleep medication, trouble sleeping, and sleep hours in the sensitivity test. In the test, each of these PSQI measures was first linked to the coupling measures and age, and then regressed out from the coupling measures to further examine the age-coupling associations studied in **Fig. 1**. Similar age-coupling association tests were applied on the whole brain volume and CSF volume, respectively. The whole brain volume was accessed by the volume number of all brain regions excluding the CSF area from the T1-weight MRI, where the CSF regions mainly from all the ventricles were extracted to quantify the CSF volume. To test whether the head motion would drive the age-coupling association, we adjusted the gBOLD-CSF coupling for the head motion from each rsfMRI acquisition, which was quantified by the session-mean framewise displacement (FD), following the previous study (Han et al., 2021b) and replicated the analysis in **Fig. 1**. The FD was calculated as the sum of the absolute value of all 6 translational and rotational realignment parameters derived from the preprocessing (Power et al., 2012). We did not use motion-censoring methods (Power et al., 2014, 2012) to avoid the influence from the cross-correlation of the concatenated timeseries of gBOLD and CSF fMRI signals.

### Sex-specific coupling changes with aging

We further compared the gBOLD–CSF coupling strength between the male and female subjects, as well as the respective trajectory of coupling changing with aging. First, we compared the coupling measures between sexes with a two-sample t-test. Second, we divided the males or females into 7 sub-groups (one group per 10 years; starting from 36∼45 yrs, i.e., 36 ≤ age < 46), respectively, and compared the coupling measures across these sub-groups.

To examine whether the menopause would affect the tendency of the coupling measure changing with aging, we separated all the females into the “post-menopause” and “other females” groups (based on the measure of “whether having no period for 12 months” in the “menstrual cycle” data), divided the “other females” into two stages of the “36∼45 yrs” and “46∼55 yrs”, and compared the coupling measures between the two stages. Similarly, we also selected the same age range/sub-groups for the post-menopausal subjects and contrasted the corresponding coupling measures between the two sub-groups, as well as compared the coupling across the “post-menopause” and “other females” subjects for each age-stage. Furthermore, we applied the same grouping metric on the “post-menopause” subjects as the entire group of females above, i.e., 7 groups with an age duration of 10 years (from 36∼45 yrs), and then replicated the analyses for whole females to observe the trajectory of the coupling changing with aging.

### Statistical analysis

The present study used the two-sample t-test for the group comparison of continuous variables, including the coupling difference of neighboring groups and that between males and females. The linear regression was applied to estimate the association between age and gBOLD–CSF coupling for the younger group and older group, respectively. The ordinal regression was used for responses with natural ordering among categories (i.e., the association between the coupling measure or age and these PSQI measures). The cross-correlation function was used to evaluate the relationship between the gBOLD signal and CSF inflow signal at different time lags. We also tested the interaction effects of age and menopause on the coupling measure. In the study, Pearson’s correlation was employed to access the inter-subject associations between different variables. A p-value less than 0.05 was considered statistical significance.

## Acknowledgements

The HCP-A dataset reported in this study was supported by grants U01AG052564 and U01AG052564-S1 and by the 14 NIH Institutes and Centers that support the NIH Blueprint for Neuroscience Research, by the McDonnell Center for Systems Neuroscience at Washington University, by the Office of the Provost at Washington University, and by the University of Minnesota Medical School. We gratefully acknowledge the efforts of all the individuals who have contributed to the project.

Data and/or research tools used in the preparation of this manuscript were obtained from the National Institute of Mental Health (NIMH) Data Archive (NDA). NDA is a collaborative informatics system created by the National Institutes of Health to provide a national resource to support and accelerate research in mental health. Dataset identifier(s): 10.15154/1528735. This manuscript reflects the views of the authors and may not reflect the opinions or views of the NIH or of the Submitters submitting original data to NDA.

## Author contributions

Conceptualization: Feng Han, Xiao Liu Methodology: Feng Han, Xiao Liu

Investigation: Feng Han, Xufu Liu, Yifan Yang, Xiao Liu Visualization: Feng Han, Yifan Yang, Xiao Liu

Funding acquisition: Xiao Liu Project administration: Xiao Liu Supervision: Xiao Liu

Writing – original draft: Feng Han, Xiao Liu

Writing – review & editing: Feng Han, Xufu Liu, Yifan Yang, Xiao Liu

## Data and Materials Availability

The multimodal data, including subject characteristics, structural MRI, and rsfMRI are all publicly available at the NDA website upon the approval of the data use application (https://nda.nih.gov/get/access-data.html). All the code used in the present study are available from the corresponding author upon request.

## Supplementary Materials

### Fig. S1 to S4 for multiple supplementary figures

**Fig. S1.**
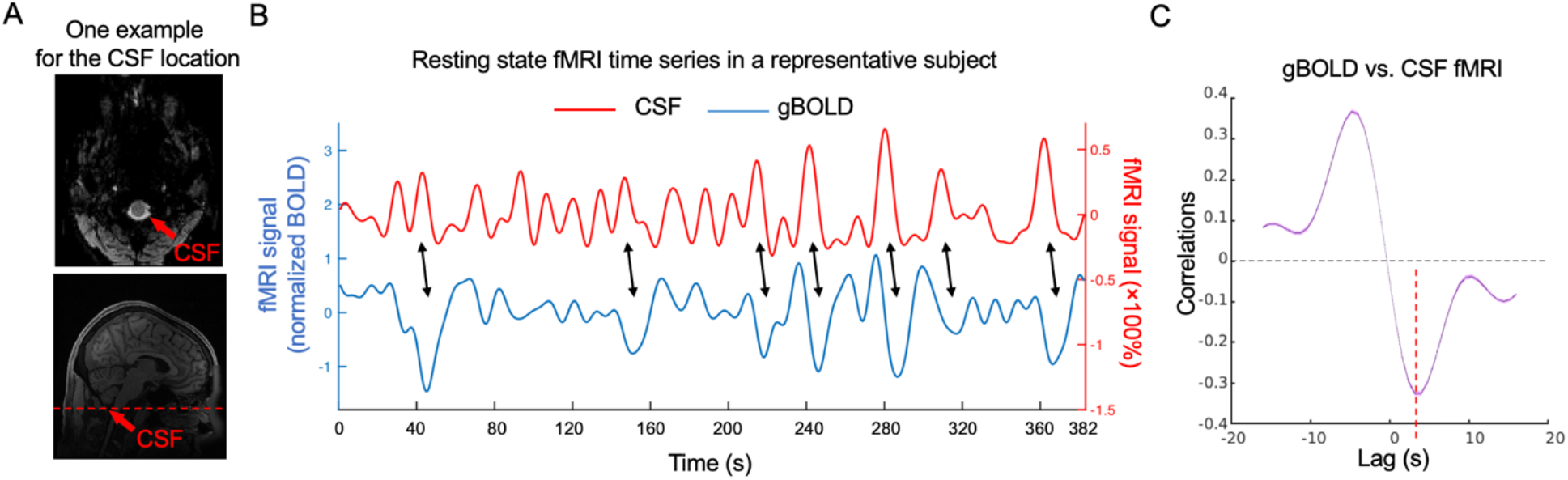
gBOLD is coupled with CSF changes in HCP-Aging data. (**A**) Top: CSF region at bottom fMRI slice shown in an example subject; Bottom: location of bottom rsfMRI slice marked in corresponding structural MRI (dashed line). (**B**) gBOLD (blue) and CSF (red) rsfMRI signals showed a coupled change (black arrows) from a representative example. (**C**) Averaged cross-correlation function between gBOLD and CSF across 719 subjects. The red dashed line marks the time lag (+3.2-sec) where the negative peak of the mean cross-correlation occurs. The shaded regions represent the area within one standard error of the mean.

**Fig. S2.**
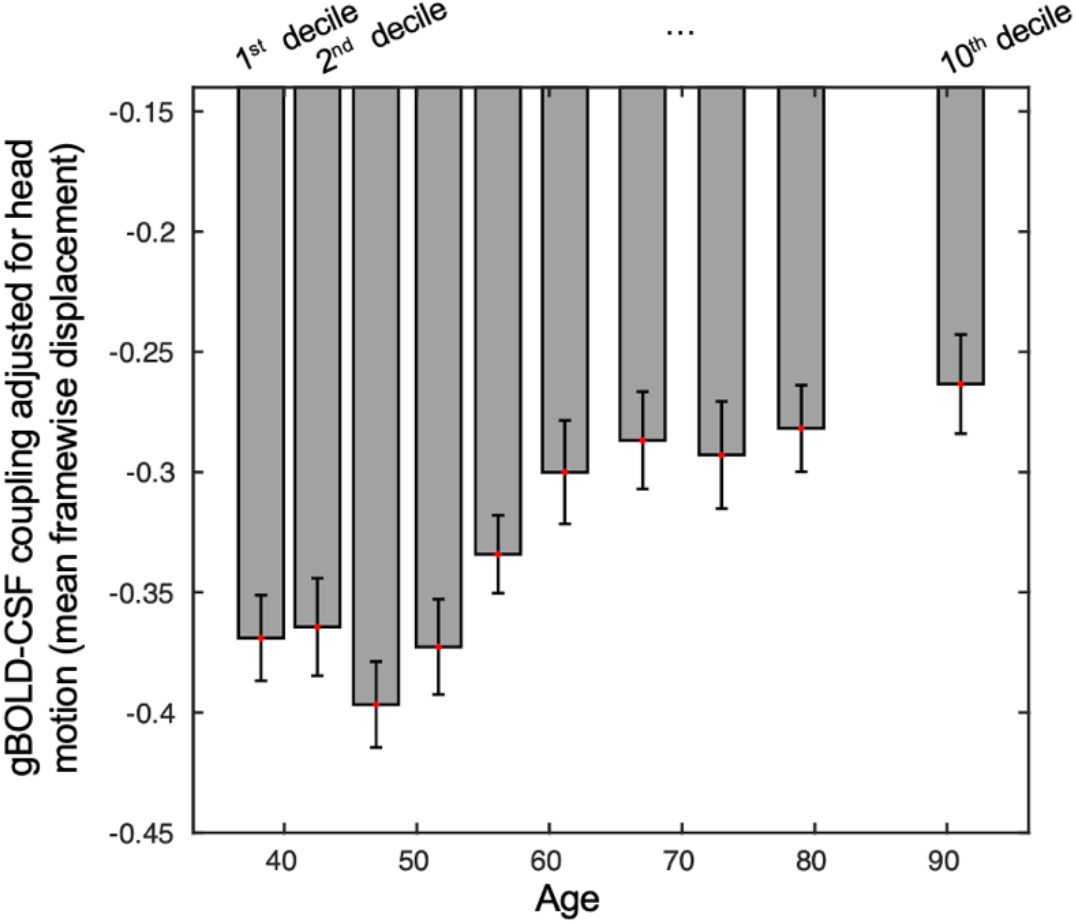
The age-coupling association was not affected by head motion during fMRI acquisition. Similar results as **Fig. 1** were found when the mean FD was regressed out from the gBOLD-CSF coupling for each rsfMRI session.

**Fig. S3.**
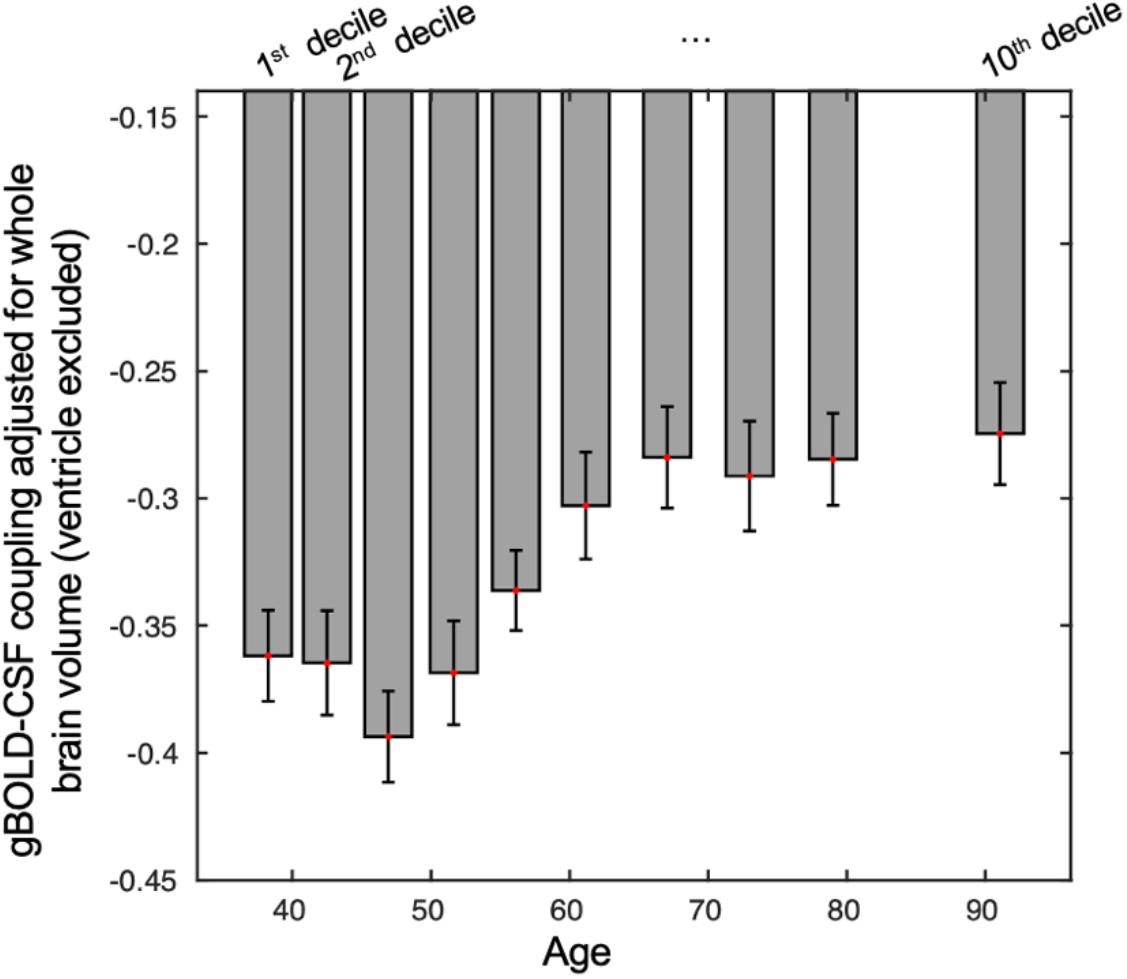
The age-coupling association was not dominant by the whole brain volume. Consistent results as **Fig. 1** were found when we regressed out the whole brain volume (ventricles excluded) from the gBOLD-CSF coupling for each rsfMRI session.

**Fig. S4.**
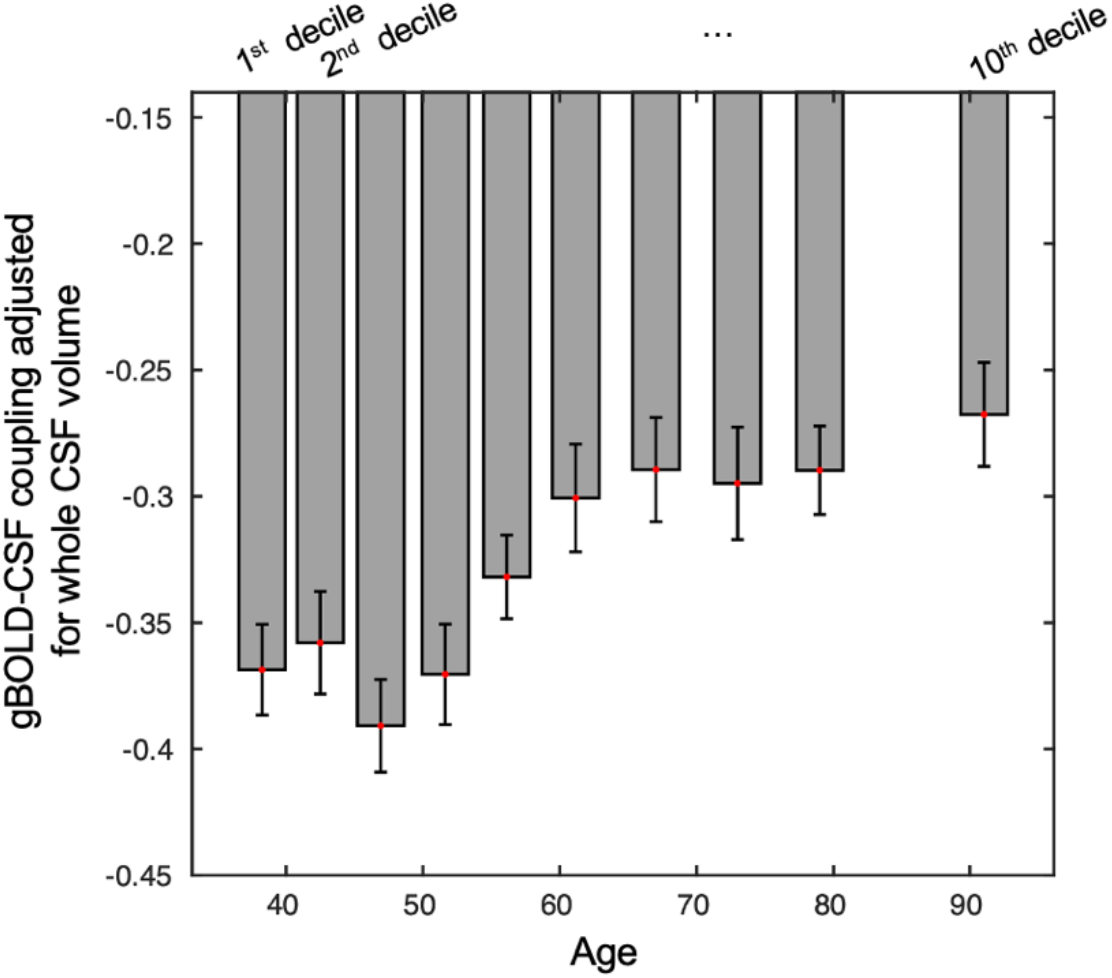
The age-coupling association was not affected by the whole CSF volume. Similar results as **Fig. 1** were found when the volume of the whole CSF area was regressed out from the gBOLD-CSF coupling for each rsfMRI session.

## References

Abbott, N.J., Pizzo, M.E., Preston, J.E., Janigro, D., Thorne, R.G., 2018. The role of brain barriers in fluid movement in the CNS: is there a ‘glymphatic’ system? Acta Neuropathol. 135, 387–407. https://doi.org/10.1007/s00401-018-1812-4

Baust, W., Bohnert, B., 1969. The regulation of heart rate during sleep. Exp. Brain Res. https://doi.org/10.1007/BF00235442

Benveniste, H., Liu, X., Koundal, S., Sanggaard, S., Lee, H., Wardlaw, J., 2019. The Glymphatic System and Waste Clearance with Brain Aging: A Review. Gerontology 65, 106–119. https://doi.org/10.1159/000490349

Birn, R.M., Diamond, J.B., Smith, M.A., Bandettini, P.A., 2006. Separating respiratory-variation-related fluctuations from neuronal-activity-related fluctuations in fMRI. Neuroimage. https://doi.org/10.1016/j.neuroimage.2006.02.048

Boland, B., Yu, W.H., Corti, O., Mollereau, B., Henriques, A., Bezard, E., Pastores, G.M., Rubinsztein, D.C., Nixon, R.A., Duchen, M.R., Mallucci, G.R., Kroemer, G., Levine, B., Eskelinen, E.-L., Mochel, F., Spedding, M., Louis, C., Martin, O.R., Millan, M.J., 2018. Promoting the clearance of neurotoxic proteins in neurodegenerative disorders of ageing. Nat. Rev. Drug Discov. 17, 660–688. https://doi.org/10.1038/nrd.2018.109

Bookheimer, S.Y., Salat, D.H., Terpstra, M., Ances, B.M., Barch, D.M., Buckner, R.L., Burgess, G.C., Curtiss, S.W., Diaz-Santos, M., Elam, J.S., Fischl, B., Greve, D.N., Hagy, H.A., Harms, M.P., Hatch, O.M., Hedden, T., Hodge, C., Japardi, K.C., Kuhn, T.P., Ly, T.K., Smith, S.M., Somerville, L.H., Uğurbil, K., van der Kouwe, A., Van Essen, D., Woods, R.P., Yacoub, E., 2019. The Lifespan Human Connectome Project in Aging: An overview. Neuroimage 185, 335–348. https://doi.org/10.1016/j.neuroimage.2018.10.009

Boudreau, P., Yeh, W.H., Dumont, G.A., Boivin, D.B., 2013. Circadian variation of heart rate variability across sleep stages. Sleep. https://doi.org/10.5665/sleep.3230

Brown, A.M.C., Gervais, N.J., 2020. Role of Ovarian Hormones in the Modulation of Sleep in Females Across the Adult Lifespan. Endocrinology 161. https://doi.org/10.1210/endocr/bqaa128

Chang, C., Cunningham, J.P., Glover, G.H., 2009. Influence of heart rate on the BOLD signal: The cardiac response function. Neuroimage. https://doi.org/10.1016/j.neuroimage.2008.09.029

Chen, R.L., Kassem, N.A., Redzic, Z.B., Chen, C.P.C., Segal, M.B., Preston, J.E., 2009. Age-related changes in choroid plexus and blood–cerebrospinal fluid barrier function in the sheep. Exp. Gerontol. 44, 289–296. https://doi.org/https://doi.org/10.1016/j.exger.2008.12.004

Cox, R.W., 1996. AFNI: Software for analysis and visualization of functional magnetic resonance neuroimages. Comput. Biomed. Res. https://doi.org/10.1006/cbmr.1996.0014

Da Mesquita, S., Louveau, A., Vaccari, A., Smirnov, I., Cornelison, R.C., Kingsmore, K.M., Contarino, C., Onengut-Gumuscu, S., Farber, E., Raper, D., Viar, K.E., Powell, R.D., Baker, W., Dabhi, N., Bai, R., Cao, R., Hu, S., Rich, S.S., Munson, J.M., Lopes, M.B., Overall, C.C., Acton, S.T., Kipnis, J., 2018. Functional aspects of meningeal lymphatics in ageing and Alzheimer’s disease. Nature 560, 185–191. https://doi.org/10.1038/s41586-018-0368-8

De Gennaro, L., Gorgoni, M., Reda, F., Lauri, G., Truglia, I., Cordone, S., Scarpelli, S., Mangiaruga, A., D’atri, A., Lacidogna, G., Ferrara, M., Marra, C., Rossini, P.M., 2017. The Fall of Sleep K-Complex in Alzheimer Disease. Sci. Rep. 7, 39688. https://doi.org/10.1038/srep39688

Douglas, N.J., White, D.P., Pickett, C.K., Weil, J. V., Zwillich, C.W., 1982. Respiration during sleep in normal man. Thorax. https://doi.org/10.1136/thx.37.11.840

Eide, P.K., Vatnehol, S.A.S., Emblem, K.E., Ringstad, G., 2018. Magnetic resonance imaging provides evidence of glymphatic drainage from human brain to cervical lymph nodes. Sci. Rep. 8, 7194. https://doi.org/10.1038/s41598-018-25666-4

Fleischman, D., Berdahl, J.P., Zaydlarova, J., Stinnett, S., Fautsch, M.P., Allingham, R.R., 2012. Cerebrospinal Fluid Pressure Decreases with Older Age. PLoS One 7, 1–9. https://doi.org/10.1371/journal.pone.0052664

Fukunaga, M., Horovitz, S.G., van Gelderen, P., de Zwart, J.A., Jansma, J.M., Ikonomidou, V.N., Chu, R., Deckers, R.H.R., Leopold, D.A., Duyn, J.H., 2006. Large-amplitude, spatially correlated fluctuations in BOLD fMRI signals during extended rest and early sleep stages. Magn. Reson. Imaging 24, 979–992. https://doi.org/10.1016/j.mri.2006.04.018

Fultz, N.E., Bonmassar, G., Setsompop, K., Stickgold, R.A., Rosen, B.R., Polimeni, J.R., Lewis, L.D., 2019. Coupled electrophysiological, hemodynamic, and cerebrospinal fluid oscillations in human sleep. Science (80-.). https://doi.org/10.1126/science.aax5440

Gao, J.H., Liu, H.L., 2012. Inflow effects on functional MRI. Neuroimage. https://doi.org/10.1016/j.neuroimage.2011.09.088

Gao, J.H., Miller, I., Lai, S., Xiong, J., Fox, P.T., 1996. Quantitative assessment of blood inflow effects in functional MRI signals. Magn. Reson. Med. https://doi.org/10.1002/mrm.1910360219

Gu, Y., Han, F., Sainburg, L.E., Liu, X., 2020. Transient Arousal Modulations Contribute to Resting-State Functional Connectivity Changes Associated with Head Motion Parameters. Cereb. Cortex. https://doi.org/10.1093/cercor/bhaa096

Gu, Y., Han, F., Sainburg, L.E., Schade, M.M., Buxton, O.M., Duyn, J.H., Liu, X., 2022. An orderly sequence of autonomic and neural events at transient arousal changes. Neuroimage. https://doi.org/10.1016/j.neuroimage.2022.119720

Gu, Y., Sainburg, L.E., Kuang, S., Han, F., Williams, J.W., Liu, Y., Zhang, N., Zhang, X., Leopold, D.A., Liu, X., 2021. Brain Activity Fluctuations Propagate as Waves Traversing the Cortical Hierarchy. Cereb. Cortex 31, 3986–4005. https://doi.org/10.1093/cercor/bhab064

Guazzi, M., Zanchetti, A., 1965. Carotid sinus and aortic reflexes in the regulation of circulation during sleep. Science (80-.). https://doi.org/10.1126/science.148.3668.397

Hablitz, L.M., Vinitsky, H.S., Sun, Q., Stæger, F.F., Sigurdsson, B., Mortensen, K.N., Lilius, T.O., Nedergaard, M., 2019. Increased glymphatic influx is correlated with high EEG delta power and low heart rate in mice under anesthesia. Sci. Adv. 5, eaav5447. https://doi.org/10.1126/sciadv.aav5447

Hachul, H., Frange, C., Bezerra, A.G., Hirotsu, C., Pires, G.N., Andersen, M.L., Bittencourt, L., Tufik, S., 2015. The effect of menopause on objective sleep parameters: Data from an epidemiologic study in São Paulo, Brazil. Maturitas 80, 170–178. https://doi.org/https://doi.org/10.1016/j.maturitas.2014.11.002

Han, F., Brown, G.L., Zhu, Y., Belkin-Rosen, A.E., Lewis, M.M., Du, G., Gu, Y., Eslinger, P.J., Mailman, R.B., Huang, X., Liu, X., 2021a. Decoupling of Global Brain Activity and Cerebrospinal Fluid Flow in Parkinson’s Disease Cognitive Decline. Mov. Disord. 36, 2066–2076. https://doi.org/10.1002/mds.28643

Han, F., Chen, J., Belkin-Rosen, A., Gu, Y., Luo, L., Buxton, O.M., Liu, X., 2021b. Reduced coupling between cerebrospinal fluid flow and global brain activity is linked to Alzheimer disease–related pathology. PLoS Biol. 19, 1–25. https://doi.org/10.1371/journal.pbio.3001233

Han, F., Liu, Xufu, Mailman, R.B., Huang, X., Liu, Xiao, 2022. Early β-amyloid accumulation and hypoconnectivity in the default mode network are related to its disengagement from global brain activity. bioRxiv. https://doi.org/10.1101/2022.07.24.501309

Harms, M.P., Somerville, L.H., Ances, B.M., Andersson, J., Barch, D.M., Bastiani, M., Bookheimer, S.Y., Brown, T.B., Buckner, R.L., Burgess, G.C., Coalson, T.S., Chappell, M.A., Dapretto, M., Douaud, G., Fischl, B., Glasser, M.F., Greve, D.N., Hodge, C., Jamison, K.W., Jbabdi, S., Kandala, S., Li, X., Mair, R.W., Mangia, S., Marcus, D., Mascali, D., Moeller, S., Nichols, T.E., Robinson, E.C., Salat, D.H., Smith, S.M., Sotiropoulos, S.N., Terpstra, M., Thomas, K.M., Tisdall, M.D., Ugurbil, K., van der Kouwe, A., Woods, R.P., Zöllei, L., Van Essen, D.C., Yacoub, E., 2018. Extending the Human Connectome Project across ages: Imaging protocols for the Lifespan Development and Aging projects. Neuroimage 183, 972–984. https://doi.org/10.1016/j.neuroimage.2018.09.060

Helakari, H., Korhonen, V., Holst, S.C., Piispala, J., Kallio, M., Väyrynen, T., Huotari, N., Raitamaa, L., Tuunanen, J., Kananen, J., Järvelä, M., Tuovinen, T., Raatikainen, V., Borchardt, V., Kinnunen, H., Nedergaard, M., Kiviniemi, V., 2022. Human NREM Sleep Promotes Brain-Wide Vasomotor and Respiratory Pulsations. J. Neurosci. JN-RM-0934-21. https://doi.org/10.1523/jneurosci.0934-21.2022

Hladky, S.B., Barrand, M.A., 2022. The glymphatic hypothesis: the theory and the evidence, Fluids and Barriers of the CNS. BioMed Central. https://doi.org/10.1186/s12987-021-00282-z

Holth, J.K., Fritschi, S.K., Wang, C., Pedersen, N.P., Cirrito, J.R., Mahan, T.E., Finn, M.B., Manis, M., Geerling, J.C., Fuller, P.M., Lucey, B.P., Holtzman, D.M., 2019. The sleep-wake cycle regulates brain interstitial fluid tau in mice and CSF tau in humans. Science (80-.). https://doi.org/10.1126/science.aav2546

Hou, Y., Dan, X., Babbar, M., Wei, Y., Hasselbalch, S.G., Croteau, D.L., Bohr, V.A., 2019. Ageing as a risk factor for neurodegenerative disease. Nat. Rev. Neurol. 15, 565–581. https://doi.org/10.1038/s41582-019-0244-7

Iliff, J.J., Chen, M.J., Plog, B.A., Zeppenfeld, D.M., Soltero, M., Yang, L., Singh, I., Deane, R., Nedergaard, M., 2014. Impairment of glymphatic pathway function promotes tau pathology after traumatic brain injury. J. Neurosci. https://doi.org/10.1523/JNEUROSCI.3020-14.2014

Iliff, J.J., Wang, M., Liao, Y., Plogg, B.A., Peng, W., Gundersen, G.A., Benveniste, H., Vates, G.E., Deane, R., Goldman, S.A., Nagelhus, E.A., Nedergaard, M., 2012. A paravascular pathway facilitates CSF flow through the brain parenchyma and the clearance of interstitial solutes, including amyloid β. Sci. Transl. Med. https://doi.org/10.1126/scitranslmed.3003748

Iliff, J.J., Wang, M., Zeppenfeld, D.M., Venkataraman, A., Plog, B.A., Liao, Y., Deane, R., Nedergaard, M., 2013. Cerebral arterial pulsation drives paravascular CSF-Interstitial fluid exchange in the murine brain. J. Neurosci. https://doi.org/10.1523/JNEUROSCI.1592-13.2013

Jack, C.R., Knopman, D.S., Jagust, W.J., Petersen, R.C., Weiner, M.W., Aisen, P.S., Shaw, L.M., Vemuri, P., Wiste, H.J., Weigand, S.D., Lesnick, T.G., Pankratz, V.S., Donohue, M.C., Trojanowski, J.Q., 2013. Tracking pathophysiological processes in Alzheimer’s disease: An updated hypothetical model of dynamic biomarkers. Lancet Neurol. 12, 207–216. https://doi.org/10.1016/S1474-4422(12)70291-0

Jessen, N.A., Munk, A.S.F., Lundgaard, I., Nedergaard, M., 2015. The Glymphatic System: A Beginner’s Guide. Neurochem. Res. https://doi.org/10.1007/s11064-015-1581-6

Ju, Y.-E.S., Ooms, S.J., Sutphen, C., Macauley, S.L., Zangrilli, M.A., Jerome, G., Fagan, A.M., Mignot, E., Zempel, J.M., Claassen, J.A.H.R., Holtzman, D.M., 2017. Slow wave sleep disruption increases cerebrospinal fluid amyloid-β levels. Brain 140, 2104–2111. https://doi.org/10.1093/brain/awx148

Kiviniemi, V., Wang, X., Korhonen, V., Keinänen, T., Tuovinen, T., Autio, J., Levan, P., Keilholz, S., Zang, Y.F., Hennig, J., Nedergaard, M., 2016. Ultra-fast magnetic resonance encephalography of physiological brain activity-Glymphatic pulsation mechanisms? J. Cereb. Blood Flow Metab. https://doi.org/10.1177/0271678X15622047

Kress, B.T., Iliff, J.J., Xia, M., Wang, M., Wei Bs, H.S., Zeppenfeld, D., Xie, L., Hongyi Kang, B.S., Xu, Q., Liew, J.A., Plog, B.A., Ding, F., PhD, R.D., Nedergaard, M., 2014. Impairment of paravascular clearance pathways in the aging brain. Ann. Neurol. 76, 845–861. https://doi.org/10.1002/ana.24271

Larson-Prior, L.J., Zempel, J.M., Nolan, T.S., Prior, F.W., Snyder, A., Raichle, M.E., 2009. Cortical network functional connectivity in the descent to sleep. Proc. Natl. Acad. Sci. U. S. A. https://doi.org/10.1073/pnas.0900924106

Levine, D.A., Gross, A.L., Briceño, E.M., Tilton, N., Giordani, B.J., Sussman, J.B., Hayward, R.A., Burke, J.F., Hingtgen, S., Elkind, M.S.V., Manly, J.J., Gottesman, R.F., Gaskin, D.J., Sidney, S., Sacco, R.L., Tom, S.E., Wright, C.B., Yaffe, K., Galecki, A.T., 2021. Sex Differences in Cognitive Decline among US Adults. JAMA Netw. Open 4, 1–13. https://doi.org/10.1001/jamanetworkopen.2021.0169

Liu, X., De Zwart, J.A., Schölvinck, M.L., Chang, C., Ye, F.Q., Leopold, D.A., Duyn, J.H., 2018. Subcortical evidence for a contribution of arousal to fMRI studies of brain activity. Nat. Commun. https://doi.org/10.1038/s41467-017-02815-3

Liu, X., Leopold, D.A., Yang, Y., 2021. Single-neuron firing cascades underlie global spontaneous brain events. Proc. Natl. Acad. Sci. U. S. A. 118, 1–10. https://doi.org/10.1073/pnas.2105395118

Liu, X., Yanagawa, T., Leopold, D.A., Chang, C., Ishida, H., Fujii, N., Duyn, J.H., 2015. Arousal transitions in sleep, wakefulness, and anesthesia are characterized by an orderly sequence of cortical events. Neuroimage. https://doi.org/10.1016/j.neuroimage.2015.04.003

Mander, B.A., Marks, S.M., Vogel, J.W., Rao, V., Lu, B., Saletin, J.M., Ancoli-Israel, S., Jagust, W.J., Walker, M.P., 2015. β-amyloid disrupts human NREM slow waves and related hippocampus-dependent memory consolidation. Nat. Neurosci. 18, 1051–1057. https://doi.org/10.1038/nn.4035

Mander, B.A., Winer, J.R., Walker, M.P., 2017. Sleep and Human Aging. Neuron 94, 19–36. https://doi.org/https://doi.org/10.1016/j.neuron.2017.02.004

Mawuenyega, K.G., Sigurdson, W., Ovod, V., Munsell, L., Kasten, T., Morris, J.C., Yarasheski, K.E., Bateman, R.J., 2010. Decreased clearance of CNS β-amyloid in Alzheimer’s disease. Science (80-.). 330, 1774. https://doi.org/10.1126/science.1197623

Mestre, H., Tithof, J., Du, T., Song, W., Peng, W., Sweeney, A.M., Olveda, G., Thomas, J.H., Nedergaard, M., Kelley, D.H., 2018. Flow of cerebrospinal fluid is driven by arterial pulsations and is reduced in hypertension. Nat. Commun. https://doi.org/10.1038/s41467-018-07318-3

Mielke, M.M., Vemuri, P., Rocca, W.A., 2014. Clinical epidemiology of Alzheimer’s disease: Assessing sex and gender differences. Clin. Epidemiol. https://doi.org/10.2147/CLEP.S37929

Nedergaard, M., Goldman, S.A., 2020. Glymphatic failure as a final common pathway to dementia. Science (80-.). 370, 50–56. https://doi.org/10.1126/science.abb8739

Nooyens, A.C.J., Wijnhoven, H.A.H., Schaap, L.S., Sialino, L.D., Kok, A.A.L., Visser, M., Verschuren, W.M.M., Picavet, H.S.J., van Oostrom, S.H., 2022. Sex Differences in Cognitive Functioning with Aging in the Netherlands. Gerontology. https://doi.org/10.1159/000520318

Olbrich, S., Mulert, C., Karch, S., Trenner, M., Leicht, G., Pogarell, O., Hegerl, U., 2009. EEG-vigilance and BOLD effect during simultaneous EEG/fMRI measurement. Neuroimage. https://doi.org/10.1016/j.neuroimage.2008.11.014

Özbay, P.S., Chang, C., Picchioni, D., Mandelkow, H., Chappel-Farley, M.G., van Gelderen, P., de Zwart, J.A., Duyn, J., 2019. Sympathetic activity contributes to the fMRI signal. Commun. Biol. https://doi.org/10.1038/s42003-019-0659-0

Özbay, P.S., Chang, C., Picchioni, D., Mandelkow, H., Moehlman, T.M., Chappel-Farley, M.G., van Gelderen, P., de Zwart, J.A., Duyn, J.H., 2018. Contribution of systemic vascular effects to fMRI activity in white matter. Neuroimage. https://doi.org/10.1016/j.neuroimage.2018.04.045

Pais-Roldán, P., Takahashi, K., Sobczak, F., Chen, Y., Zhao, X., Zeng, H., Jiang, Y., Yu, X., 2020. Indexing brain state-dependent pupil dynamics with simultaneous fMRI and optical fiber calcium recording. Proc. Natl. Acad. Sci. U. S. A. 117, 6875–6882. https://doi.org/10.1073/pnas.1909937117

Palmqvist, S., Schöll, M., Strandberg, O., Mattsson, N., Stomrud, E., Zetterberg, H., Blennow, K., Landau, S., Jagust, W., Hansson, O., 2017. Earliest accumulation of β-amyloid occurs within the default-mode network and concurrently affects brain connectivity. Nat. Commun. https://doi.org/10.1038/s41467-017-01150-x

Peng, W., Achariyar, T.M., Li, B., Liao, Y., Mestre, H., Hitomi, E., Regan, S., Kasper, T., Peng, S., Ding, F., Benveniste, H., Nedergaard, M., Deane, R., 2016. Suppression of glymphatic fluid transport in a mouse model of Alzheimer’s disease. Neurobiol. Dis. 93, 215–225. https://doi.org/https://doi.org/10.1016/j.nbd.2016.05.015

Power, J.D., Barnes, K.A., Snyder, A.Z., Schlaggar, B.L., Petersen, S.E., 2012. Spurious but systematic correlations in functional connectivity MRI networks arise from subject motion. Neuroimage. https://doi.org/10.1016/j.neuroimage.2011.10.018

Power, J.D., Mitra, A., Laumann, T.O., Snyder, A.Z., Schlaggar, B.L., Petersen, S.E., 2014. Methods to detect, characterize, and remove motion artifact in resting state fMRI. Neuroimage 84, 320–341. https://doi.org/10.1016/j.neuroimage.2013.08.048

Power, J.D., Plitt, M., Gotts, S.J., Kundu, P., Voon, V., Bandettini, P.A., Martin, A., 2018. Ridding fMRI data of motion-related influences: Removal of signals with distinct spatial and physical bases in multiecho data. Proc. Natl. Acad. Sci. U. S. A. https://doi.org/10.1073/pnas.1720985115

Qiu, C., Kivipelto, M., von Strauss, E., 2009. Epidemiology of Alzheimer’s disease: occurrence, determinants, and strategies toward intervention. Dialogues Clin. Neurosci. 11, 111–128. https://doi.org/10.31887/DCNS.2009.11.2/cqiu

Raut, R. V., Snyder, A.Z., Mitra, A., Yellin, D., Fujii, N., Malach, R., Raichle, M.E., 2021. Global waves synchronize the brain’s functional systems with fluctuating arousal. Sci. Adv. 7, 1–16. https://doi.org/10.1126/sciadv.abf2709

Redline, S., Kirchner, H.L., Quan, S.F., Gottlieb, D.J., Kapur, V., Newman, A., 2004. The Effects of Age, Sex, Ethnicity, and Sleep-Disordered Breathing on Sleep Architecture. Arch. Intern. Med. 164, 406–418. https://doi.org/10.1001/archinte.164.4.406

Smith, S.M., Jenkinson, M., Woolrich, M.W., Beckmann, C.F., Behrens, T.E.J., Johansen-Berg, H., Bannister, P.R., De Luca, M., Drobnjak, I., Flitney, D.E., Niazy, R.K., Saunders, J., Vickers, J., Zhang, Y., De Stefano, N., Brady, J.M., Matthews, P.M., 2004. Advances in functional and structural MR image analysis and implementation as FSL. Neuroimage 23, S208–S219. https://doi.org/https://doi.org/10.1016/j.neuroimage.2004.07.051

Snyder, F., Hobson, J.A., Morrison, D.F., Goldfrank, F., 1964. Changes in respiration, heart rate, and systolic blood pressure in human sleep. J. Appl. Physiol. https://doi.org/10.1152/jappl.1964.19.3.417

Sperling, R., Mormino, E., Johnson, K., 2014. The evolution of preclinical Alzheimer’s disease: Implications for prevention trials. Neuron. https://doi.org/10.1016/j.neuron.2014.10.038

Tarasoff-Conway, J.M., Carare, R.O., Osorio, R.S., Glodzik, L., Butler, T., Fieremans, E., Axel, L., Rusinek, H., Nicholson, C., Zlokovic, B. V., Frangione, B., Blennow, K., Ménard, J., Zetterberg, H., Wisniewski, T., De Leon, M.J., 2015. Clearance systems in the brain-Implications for Alzheimer disease. Nat. Rev. Neurol. https://doi.org/10.1038/nrneurol.2015.119

Thompson, G.J., Pan, W.-J., Magnuson, M.E., Jaeger, D., Keilholz, S.D., 2014. Quasi-periodic patterns (QPP): Large-scale dynamics in resting state fMRI that correlate with local infraslow electrical activity. Neuroimage 84, 1018–1031. https://doi.org/https://doi.org/10.1016/j.neuroimage.2013.09.029

Turchi, J., Chang, C., Ye, F.Q., Russ, B.E., Yu, D.K., Cortes, C.R., Monosov, I.E., Duyn, J.H., Leopold, D.A., 2018. The Basal Forebrain Regulates Global Resting-State fMRI Fluctuations. Neuron. https://doi.org/10.1016/j.neuron.2018.01.032

Turner, K.L., Gheres, K.W., Drew, P.J., 2022. Relating Pupil Diameter and Blinking to Cortical Activity and Hemodynamics across Arousal States. Journal of Neuroscience. https://doi.org/10.1523/JNEUROSCI.1244-22.2022

Turner, K.L., Gheres, K.W., Proctor, E.A., Drew, P.J., 2020. Neurovascular coupling and bilateral connectivity during NREM and REM sleep. Elife 9, e62071. https://doi.org/10.7554/eLife.62071

van Veluw, S.J., Hou, S.S., Calvo-Rodriguez, M., Arbel-Ornath, M., Snyder, A.C., Frosch, M.P., Greenberg, S.M., Bacskai, B.J., 2020. Vasomotion as a Driving Force for Paravascular Clearance in the Awake Mouse Brain. Neuron. https://doi.org/10.1016/j.neuron.2019.10.033

Varga, A.W., Wohlleber, M.E., Giménez, S., Romero, S., Alonso, J.F., Ducca, E.L., Kam, K., Lewis, C., Tanzi, E.B., Tweardy, S., Kishi, A., Parekh, A., Fischer, E., Gumb, T., Alcolea, D., Fortea, J., Lleó, A., Blennow, K., Zetterberg, H., Mosconi, L., Glodzik, L., Pirraglia, E., Burschtin, O.E., de Leon, M.J., Rapoport, D.M., Lu, S., Ayappa, I., Osorio, R.S., 2016. Reduced Slow-Wave Sleep Is Associated with High Cerebrospinal Fluid Aβ42 Levels in Cognitively Normal Elderly. Sleep 39, 2041–2048. https://doi.org/10.5665/sleep.6240

Wang, M., He, Y., Sejnowski, T.J., Yu, X., 2018. Brain-state dependent astrocytic Ca2+ signals are coupled to both positive and negative BOLD-fMRI signals. Proc. Natl. Acad. Sci. U. S. A. 115, E1647–E1656. https://doi.org/10.1073/pnas.1711692115

Winer, J.R., Mander, B.A., Kumar, S., Reed, M., Baker, S.L., Jagust, W.J., Walker, M.P., 2020. Sleep Disturbance Forecasts β-Amyloid Accumulation across Subsequent Years. Curr. Biol. 30, 4291–4298.e3. https://doi.org/10.1016/j.cub.2020.08.017

Xie, L., Kang, H., Xu, Q., Chen, M.J., Liao, Y., Thiyagarajan, M., O’Donnell, J., Christensen, D.J., Nicholson, C., Iliff, J.J., Takano, T., Deane, R., Nedergaard, M., 2013. Sleep drives metabolite clearance from the adult brain. Science (80-.). https://doi.org/10.1126/science.1241224

Yamada, S., Miyazaki, M., Yamashita, Y., Ouyang, C., Yui, M., Nakahashi, M., Shimizu, S., Aoki, I., Morohoshi, Y., McComb, J.G., 2013. Influence of respiration on cerebrospinal fluid movement using magnetic resonance spin labeling. Fluids Barriers CNS. https://doi.org/10.1186/2045-8118-10-36

Zhou, Y., Cai, J., Zhang, W., Gong, X., Yan, S., Zhang, K., Luo, Z., Sun, J., Jiang, Q., Lou, M., 2020. Impairment of the Glymphatic Pathway and Putative Meningeal Lymphatic Vessels in the Aging Human. Ann. Neurol. 87, 357–369. https://doi.org/https://doi.org/10.1002/ana.25670

Zieman, S.J., Melenovsky, V., Kass, D.A., 2005. Mechanisms, Pathophysiology, and Therapy of Arterial Stiffness. Arterioscler. Thromb. Vasc. Biol. 25, 932–943. https://doi.org/10.1161/01.ATV.0000160548.78317.29

Zou, W., Pu, T., Feng, W., Lu, M., Zheng, Y., Du, R., Xiao, M., Hu, G., 2019. Blocking meningeal lymphatic drainage aggravates Parkinson’s disease-like pathology in mice overexpressing mutated α-synuclein. Transl. Neurodegener. 8, 7. https://doi.org/10.1186/s40035-019-0147-y

